# Applicability of Castration Model in Sex Difference Studies: Insights from Metabolome and Transcriptome Analyses

**DOI:** 10.1101/2023.12.27.573488

**Authors:** Jianjun Jiang, Na Ge, Yuzhi Wang, Juntao Qi, Guibiao Wen, Xiufen Gu, Xuewen Yu, Muming Shao, Yueming Luo, Kangshuyun Gu, Feng Lin, Shudong Yang, Wei Wei

**Author notes:** Corresponding authors: Jianjun Jiang, Ph.D., Shudong Yang Wei Wei, Ph.D.

## Abstract

**Background:** Sex, as a critical biological variable, has historically been underappreciated, despite the pervasive influence of sexual dimorphism across physiological and pathological processes. A significant obstacle to advancing sex-biased biological research is the absence of an effective animal model. In recent years, castration has emerged as a potential model for elucidating sex-based differences in the context of healthy aging, where it has been shown to equalize lifespan and growth trajectories in genetically diverse mice. However, the molecular shifts induced by castration in common laboratory models, such as C57BL/6 mice, and the broader applicability of this model to other sex-related biological contexts remain largely unexplored.

**Methods:** We employed multi-omics and observational analyses to investigate the molecular changes associated with sex and sex hormones following castration. We analyzed serum, kidney, and liver samples from 12-week-old and 18-month-old castrated male C57BL/6 mice, alongside intact male and female counterparts. The castration model was further applied to assess differences in cisplatin-induced toxicity and age-related cognitive decline in comparison to unaltered controls.

**Results:** LC-MS/MS metabolomics revealed that castrated males exhibited substantial alterations in steroid hormone levels and increased concentrations of antioxidant compounds, such as taurine, despite identical diets. Integrated metabolome-transcriptome analysis confirmed distinct patterns of lipid peroxidation and oxidative stress across sham-operated female, male, and castrated male mice. Histopathological evaluations following cisplatin treatment and aging-related behavioral tests demonstrated the model’s utility in investigating sex-dependent drug toxicity and cognitive decline. These findings underscored the critical role of sex hormones in modulating both toxicity defense mechanisms and cognitive performance.

**Conclusion:** This study provides a systematic multi-omics spectrum on the castration model and demonstrates its capacity to feminize metabolic and transcriptomic profiles, establishing it as a valuable tool for exploring sex hormone-driven biological differences. Our findings lay the groundwork for further mechanistic studies and broaden the potential applications of the castration model in diverse biomedical research domains.

## INTRODUCTION

Sex is an fundamental variable that shapes physical, psychological, cognitive, and behavioral traits^1^. It exerts a significant influence on development, reproduction, and pathogenesis^2^. For instance, the incidence and survival rates of various cancers, along with their clinical phenotypes and treatment responses, exhibit marked sex-related differences^3, 4^. Physiological and immunological disparities between males and females further affect drug metabolism and toxicity during cancer therapies^5^. In aging studies, it has been well-established that women generally live longer than men, with consistent lower biological ages assessed by molecular biomarkers. However, there is a paradox that women experience a higher incidence of chronic, age-related degenerative diseases compared to men, despite their longer lifespan^6, 7^. Although awareness of sex-related differences in health and disease is rising, the molecular underpinnings of these differences remain underexplored.

A major challenge in studying sex differences lies in the lack of reliable animal models that can disentangle the complex effects of both genetic variability (such as the influence of X and Y chromosomes) and the actions of sex hormones^8^. Traditional approaches to compare males and females often fail to isolate these variables, leaving key mechanisms obscured. In recent years, castration has emerged as a potent model for elucidating the sex-based differences, particularly in healthy aging. Accumulating evidence suggests that castration can extend the lifespan across multiple mammal species. Notably, historical records indicate that Korean eunuchs had a lifespan extension of 14.4–19.1 years compared to their non-castrated counterparts of similar socio-economic status^9^. Similarly, in veterinary and livestock practices, spaying and castration are recommended for disease prevention and lifespan extension^10^. Despite the widespread applications of castration in these fields, comprehensive studies linking castration to broader biological and molecular sex differences remain sparse.

Recent studies have begun to shed light on the biological impact of castration. In 2021, Sugrue et al. provided the first evidence that castration slows the intrinsic aging process in sheep, as evidenced by a slower DNA methylation clock^11^. More recently, Jiang et al. demonstrated that prepubertal castration abolished the differences in lifespan and growth trajectories between male and female UM-HET3 mice, a genetically diverse strain^12^. While these findings offer valuable insights into the role of castration in aging, the full extent of castration’s role in other sex-relevant differences are largely unknown.

In this study, we present the first integrated multi-omics and behavioral analyses of castrated mice and their non-altered counterparts. Given that genome sequences retain the same pre-and post-castration, we hypothesize that the alterations in hormone spectrum and subsequent changes in the metabolome, transcriptome, and epigenome induced by castration could be pivotal in regulating sex dimorphism. To this end, we employed mass spectrometry-based metabolomics and deep RNA sequencing (RNA-seq) to extensively investigate the multifaceted aspects of sex-related differences associated with castration. Our results reveal several mechanistic insights, laying the groundwork for further detailed investigations and highlighting the potential of castration models as a robust tool for studying sex differences across diverse biological contexts.

## RESULTS

### Castration slowed body weight acquisition and affected the secretion of steroid hormone spectrum

To elucidate the molecular changes caused by castration, we employed multi-omics approaches to resolve the potential roles at both molecular and phenotypic levels (**Figure 1A**). Young adult C57BL/6 mice were castrated at 8 weeks old (casM) and exhibited slower gaining of body weight post-castration, but attained similar body weights as the sham males (shamM) by 60 weeks (**Figure 1B**). This pattern align with findings in prepubertal castration models using UM-HET3 mice (**Figure 1B**). At 27-30 weeks of age, we assessed the food intake among the three groups of mice. The casM group exhibited reduced food consumption compared to both shamM and sham female mice (shamF) (**Figure 1C**). This spontaneous reduction in food intake in casM may contribute to the slower body weight acquisition post-castration, and is particularly intriguing given the well-established benefits of dietary restriction for promoting healthy aging and longevity. Considering the potential impact of testosterone deprivation due to castration on the development of young adult mice, we measured their body length and found no significant difference between casM and sham (**Figure 1D**). Similarly, rectal temperature measurements revealed no significant differences among the three groups (**Figure 1E**). Notably, hair graying, a common aging marker in both mice and humans^13^, was significantly delayed in casM compared to shamM at 13 months, indicating a more youthful phenotype in the castrated group (**Supplementary Figure 1A**).

**Figure 1.**
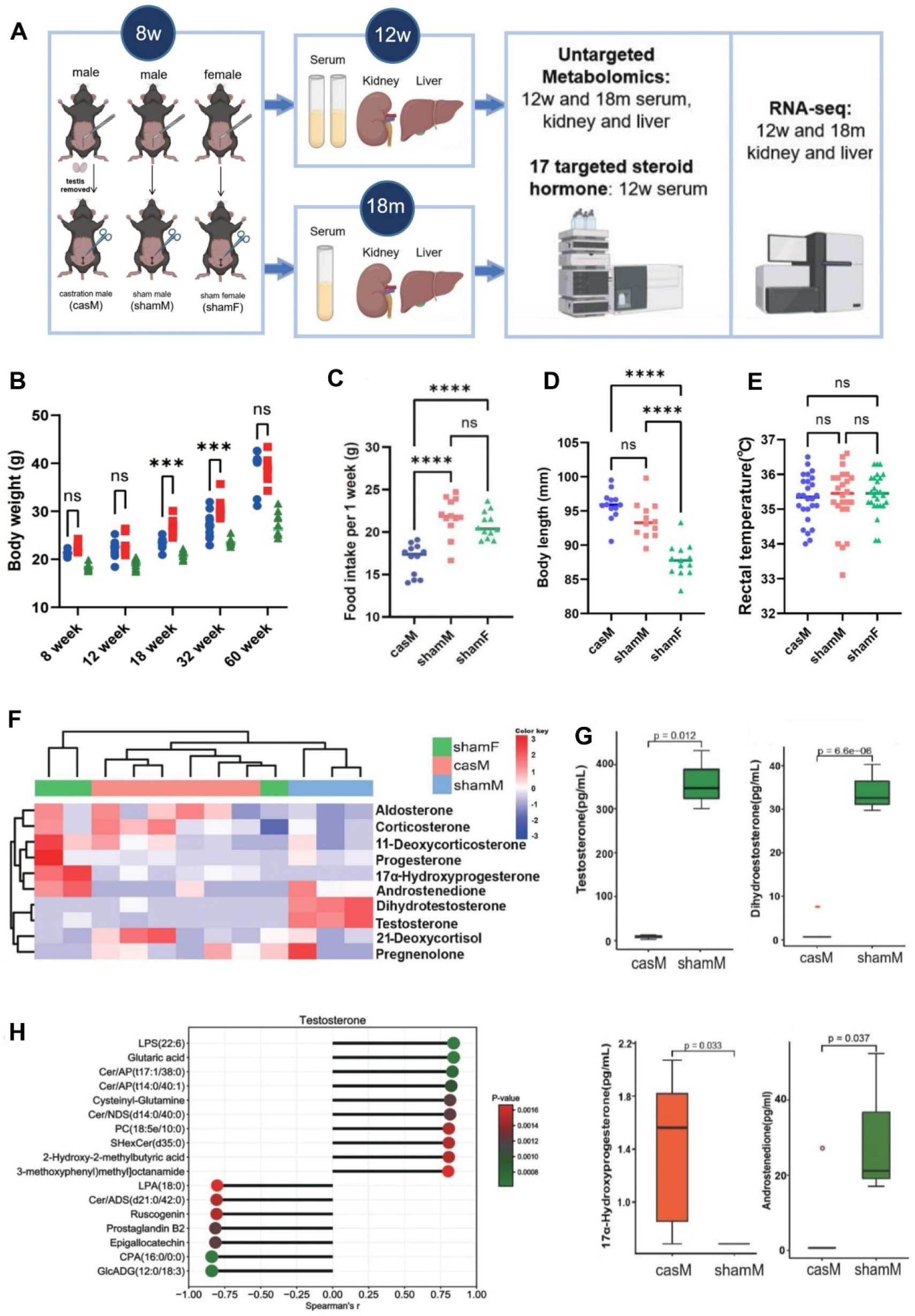
Overall study design as well as phenotypic tests and steroid hormone spectrum of castration mice. (**A**) Graphic illustration of the overall experimental design. (**B**) Body weight of casM, shamM and shamF mice from 8 to 60 weeks (n = 13, 12, 13 for casM, shamM, and shamF, respectively; unpaired students’ t-test,*\*\*\*p* < 0.001, ns: not significant) (**C**) Significant less food intake was observed in casM group by One-Way ANOVA, *\*\*\*\*p* < 0.0001 (**D-E**) non-significant difference among the 3 groups of mice was observed in (**D**) body length at 27 weeks old; (**E**) rectal temperature at 30 and 32 weeks old; (**F**) Hierarchical cluster heatmap of z-score standardized expression levels of 10 detectable steroid hormones among shamM, casM and shamF; (**G**) Comparisons of the quantitative levels of testosterone, dihydroestosterone, 17-hydroxyprogesterone and androstenedione between shamM and casM (unpaired students’ t-test, *p-*values were shown on top); (**H**) List of compounds highly correlated with TS (Spearman’s |r| ≥ 0.8, *p* ≤ 0.001). Median line was shown for G.

Hormones, serving as vital chemical messengers, travel through from bloodstream to tissues for coordinating a range of biological functions, with their dysregulation impacting development^14^, aging^15^ and disease progression^16^. The effect of surgical castration, specifically the removal of the testes, leads to a marked reduction in testosterone (TS) levels. However, the broader hormone changes resulting from castration remain underexplored. Here, we employed ultra-high-performance liquid chromatography coupled with multiple reaction monitoring tandem mass spectrometry (UHPLC-MRM-MS/MS) to analyze 17 targeted steroid hormones (SHs) in the serum of casM, shamM and shamF mice, at four weeks post-operation.

Given the limited volume of mouse serum available for SH measurement, we could only detect and compare 10 SH types across three groups. Notably, castration significantly altered SH secretion in casM (**Figure 1F**). As expected, testosterone and its more potent form, dihydrotestosterone, were remarkably reduced in CasM, to the point of being barely detectable (**Figure 1G**). Intriguingly, CasM mice exhibited a significant higher level of 17α-Hydroxyprogesterone (17α-OHP) (**Figure 1G**) and lower androstenedione (A4) compared to shamM (p < 0.05) (**Figure 1G**). These hormones, precursors to TS, might be part of a compensatory mechanism in response to TS deprivation ^17^. In addition, we observed slight increases in aldosterone (ALD) and corticosterone (CORT) in casM, with p-values of 0.11 and 0.083, respectively (**Supplementary Figure 2**). ALD, a mineralocorticoid, and CORT, a glucocorticoid, are primarily produced by the adrenal cortex. Their upregulation may suggest an activated stress response and changes in mineral and glucose homeostasis consequent to TS deprivation.

To elucidate metabolites associated with TS, we conducted an untargeted metabolomics analysis by LC-MS/MS on serum samples from three groups of mice, four weeks post-operation. A total of 1495 compounds with MS^2^ scores ≥ 0.3 were identified. Notably, coexpression analysis revealed that 17 of these compounds, which include 11 bioactive lipids, showed a significant correlation with TS levels (|r| ≥ 0.8, p ≤ 0.001, **Figure 1H**), These 11 lipids encompass a range of molecules such as prostaglandin B2 (PGD2), a specific phosphatidylcholine (PC) variant, lysophosphatidylserine (LPS), lysophosphatidic acid (LPA), cyclic PA (CPA), glucuronosyldiacylglycerol (GlcADG), sulfurHexosylceramide (SHexCer), and four types of ceramide fatty acids (Cer) (**Supplementary Table 1**). While further investigation is required to solidify these findings, our coexpression analysis suggests that TS deprivation may significantly influence peripheral lipid metabolism, particularly impacting ceramide and lysophosphatide pathways.

### Characterization of sex-related serum metabolomic features

To assess the serum metabolomic profile differences among shamM, shamF, and CasM mice at 12 weeks old (12w, 4 weeks post sham/castration procedures), principal component analysis (PCA) was performed using data from 1495 compounds identified by LC-MS/MS (**Figure 2A**). Lipid and lipid-like molecules constituted the largest category (∼33%) of these compounds (**Figure 2B**). A comparison of shamM and shamF groups to explore sex-specific differences in serum metabolic activity revealed 255 significantly altered metabolites (SAMs) were identified (**Supplementary Table 2**), with the top SAMs (ranked by fold changes) displayed in **Supplementary Figure 3A.** Despite all mice receiving the same diet, we observed notably higher levels of antioxidants and anti-inflammatory compounds in shamF compared to shamM. These compounds, likely derive from food intake, included carnosic acid^18^, arachidonic acid^19^, choline^20^, nicotinamide^21^, D-pantothenic acid^22^, gamma-crocetin^23^, ruscogenin^24^, 4-hydroxycinnamic acid^25^, and oleamide^26, 27^ (**Figure 2C**). Conversely, shamM mice exhibited elevated levels of metabolites associated with resistance to high-fat diet-related diseases, such as 5-HEPE^28^ and 12-HEPE^29^, and a higher level of protective L-Anserine^30^ than shamF (**Figure 2C**). Consistently, obesity-related compounds such as LysoPE(0:0/20:3(11Z,14Z,17Z)/0:0) (LysoPE)^31^ and 7−Ketocholesterol (7-KC)^32^ were found to be lower in male (**Figure 2C**), offering insights into why obesity is more prevalent in women than men^33^.

**Figure 2.**
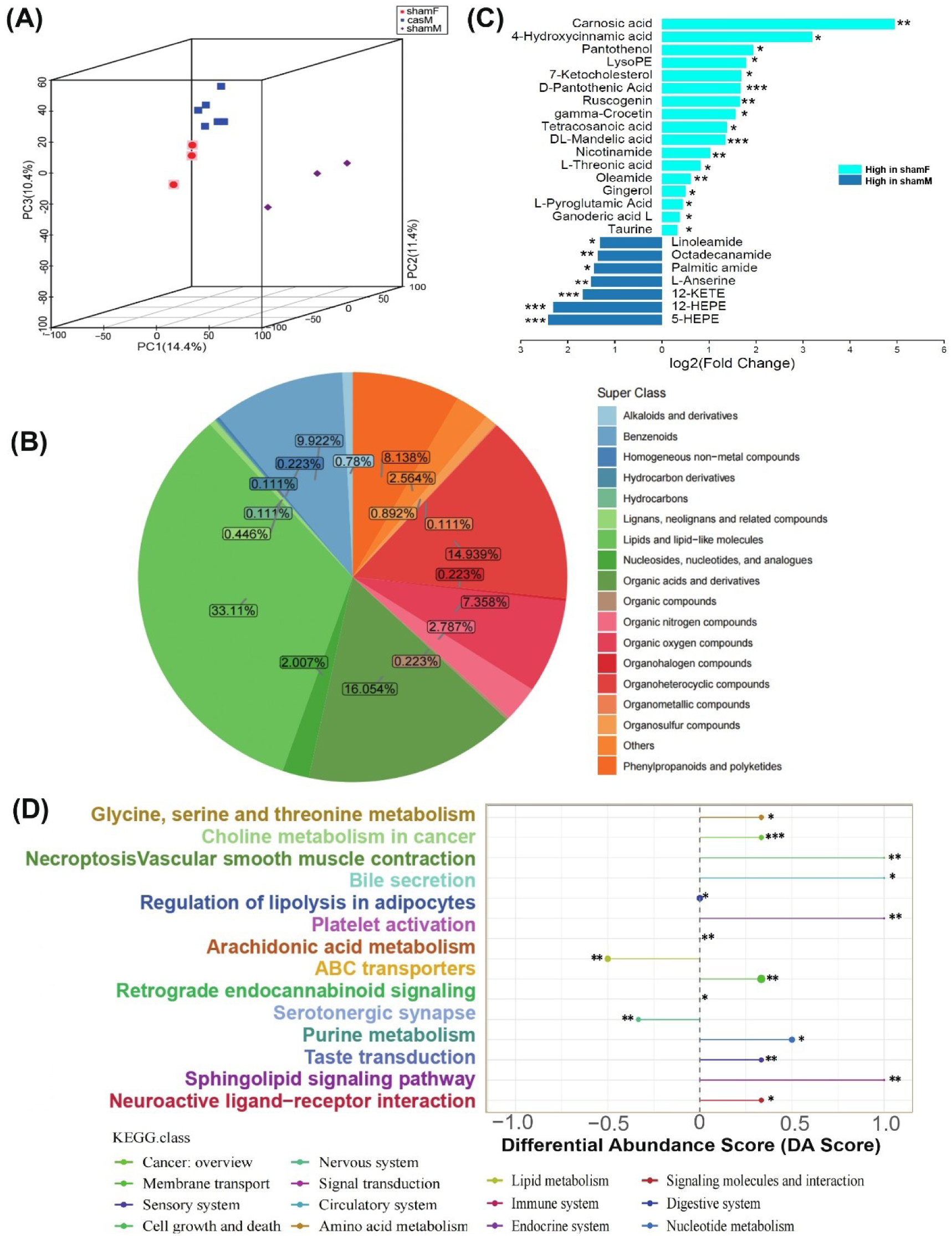
Characterizations of the 12w serum metabolomics. (**A**) PCA clustering of the serum untargeted metabolomics among casM, shamM and shamF; (**B**) Categories of the 1495 12w serum metabolites with high quality MS^2^ annotations; (**C**) Representative SAMs between shamM and shamF that related with aging and healthy aging (unpaired students’ t-test, \**p* < 0.05, \*\**p* < 0.01, \*\*\**p* < 0.001); (**D**) Differential abundance (DA) scores were calculated for the significant enrichment terms between shamM and shamF (\**p* < 0.05, \*\**p* < 0.01, \*\*\**p* < 0.001).

Differential abundance (DS) score of the 255 SAMs was calculated to inform the overall changes of each pathway (Figure 3D). These SAMs were found to be highly enriched in lipid metabolism related pathways such as “sphingolipid signaling pathway”, “regulation of lipolysis in adipocytes” and “arachidonic acid metabolism”, suggesting a sex dimorphism in lipid metabolism. Additionally, a serum untargeted metabolomic analysis in the same groups at 18 months old identified 868 compounds, with “lipid and lipid-like molecules” again being the predominant category” (**Supplementary Figure 3B**). PCA analysis suggested that the metabolic profile of 18-month-old casM mice closely resembled that of shamF (**Supplementary Figure 3C**). Similar to the younger mice, older shamF demonstrated higher levels of protective compounds derived from the same diet compared to shamM **(Supplementary Figure 3D and Supplementary Table 3**), suggesting a potentially stronger ability in females to generate healthy compounds from the same chow.

**Figure 3.**
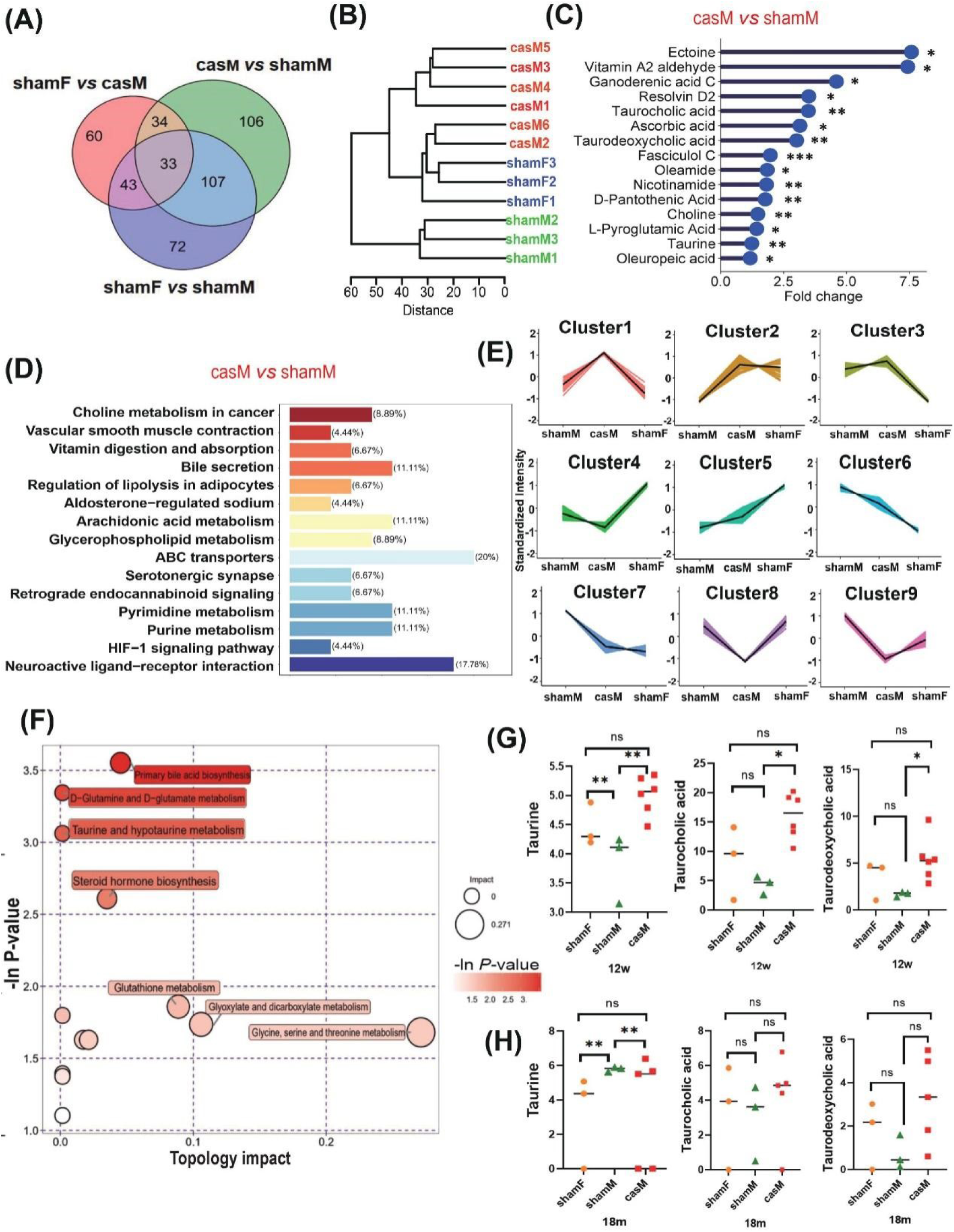
Identification of castration-specific metabolomic features in 12w mice. (**A**) Venny diagram of the serum SAMs identified from “shamF *vs* ShamM”, “shamF *vs* casM” and “casM *vs* ShamM”; (**B**) Hierarchical clustering tree of different mouse groups using levels of compounds as input; (**C**) Representative food derived SAMs between casM and shamM (unpaired students’ t-test, \**p* < 0.05, \*\**p* < 0.01, \*\*\**p* < 0.001); (D) Barplot of the KEGG enrichment terms using SAMs of “casM *vs* ShamM”, percentage of hit compounds to total SAMs was calculated for each term; (**E**) Expression patterns of merged SAMs among casM, shamM and shamF analyzed using mfuzzing clustering. Clusters 1 and 8 distinctively represent metabolites that are, respectively, highly and lowly detected specifically in casM. (**F**) Significantly enriched KEGG terms of metabolites highly expressed metabolites in casM (cluster 1 in **E**) calculated by topology-based pathway enrichment with topology impact shown in x-axis; (**G-H**) Comparisons of taurine, taurocholic acid and taurodeoxycholic acid among 12w (**G**) and 18m (**H**) shamF, shamM and casM (One-Way ANOVA, \**p* < 0.05, \*\**p* < 0.01, ns: not significant). For G and H, Tukey correction was used for correction of multiple comparisons and median lines were shown.

### Metabolomic feminization and taurine enrichment in castrated male mice

In our analysis of 12-week-old mice serum, a comparative study between casM and shamF mice identified 170 SAMs. In contrast, comparisons of “casM *vs* shamM” and “shamF *vs* shamM” revealed 280 and 255 SAMs, respectively (**Figure 3A**, **Supplementary Table 4**). Hierarchical Clustering of the 1495 identified compound demonstrated a closer metabolic profile between casM and shamF than between casM and shamM (**Figure 3B**). These findings, achieved without altering genetic background, suggest that castration feminizes the metabolomic traits of male mice. When ranking these SAMs by fold change (FC), notable differences were observed between shamM and casM. The most significantly upregulated metabolite in casM was the unsaturated fatty acid, PI (16:1/18:2), with log2FC of 13.14, while the most downregulated was guanosine, with a log_2_FC of **-**6.79. CasM mice exhibited elevated levels of various metabolites associated with healthy aging, anti-skin aging, and anti-inflammation, such as ectoine^34^, resolvin D2^35^, oleamide^26, 27^, taurine^36^, taurocholic acid^37^, taurodeoxycholic acid^38^, nicotinamide^39^, ascorbic acid (vitamin C) ^40^, D-pantothenic acid (vitamin B5)^41, 42^ and choline^43^ (**Figure 3C**). Similar to shamF, casM mice also showed an enrichment of anti-oxidative metabolites likely derived from plant-based foods, including ganoderenic acid C^44^, oleuropeic acid^45^, fasciculol C^46^, graveoline^47^ (**Figure 3C**).

Enrichment analysis of the 280 SAMs from the “casM *vs* shamM” comparison categorized them into pathways such as “bile secretion”, “vitamin digestion and absorption”, “HIF-1 signaling pathway”, “pyrimidine metabolism”, various fatty acid metabolism pathways including “arachidonic acid metabolism” and “regulation of lipolysis in adipocytes” (**Figure 3D**). Interestingly, lipid metabolism-related pathways were consistently enriched in both the “casM *vs* shamM” and “shamF *vs* shamM” comparisons, indicating a potential sensitivity of lipid metabolism to the levels of SHs.

To investigate the metabolomic changes particularly derived from castration, union SAMs among 12w-old casM, shamM and shamF groups were further classified by c-means fuzzy clustering (**Figure 3E**, **Supplementary Table 5**). This analysis revealed that cluster1 metabolites were notably elevated in casM compared to both shamM and shamF. Topology-based pathway enrichment analysis indicated that cluster1 metabolites were significantly (p < 0.05) enriched in pathways like “primary bile acid biosynthesis”, “D-glutamine and D-glutamate metabolism”, “taurine and hypotaurine metabolism” among others (**Figure 3F**). Notably, taurine levels were higher in shamF than in shamM, and castration further augmented taurine levels along with two key components of the taurine and hypotaurine metabolism pathway: taurodeoxycholic acid (TDCA) and taurocholic acid (TCA) (**Figure 3G**). However, in 18m-old mice, a higher level of taurine with no significant changes of TDAC and TCA were observed in shamM compared to shamF and casM, (**Figure 3H**), suggesting an age-related sex dimorphism of taurine concentrations in mice serum. Bile acids (BA) are crucial for maintaining lipid homeostasis and have been recognized as significant signaling molecules during aging process^47^. Given that the SAMs from both “casM *vs* shamM” and “shamF *vs* shamM” comparisons were significantly enriched in lipid-related metabolic pathways, and the primary function of taurine is the conjugation of cholesterol into bile acids^48^, our findings suggest a potential link between taurine and the sex and/or SH-related differences in lipid metabolism.

### Co-expression analysis across serum, livers and kidneys identifies potential taurine-related regulators

Kidney and liver are two major organs of organisms, the sex differences of renal and hepatic function have been aware over years^49^. To elucidate the sex and age-related regulatory networks in kidney and liver functions, we analyzed metabolome and transcriptome of renal cortex and left liver lobe from three groups of mice at 12 week (12w) and 18 months (18m) old. We detected an average of 899 compounds across both tissues, with 121 compounds common among kidneys, livers and serum samples in both age groups of sham, shamF, and casM mice (**Figure 4A** and **Supplementary Table 6**). Coexpression analysis of these 121 commonly detected metabolites (CDMs) revealed 4 major clusters (**Figure 4B**). Metabolite Set Enrichment Analysis (MSEA) indicated that Clusters 1, 2 and 3 are primarily involved in “Bile Acid Biosynthesis” and “Glycine and Serine Metabolism” pathways (**Supplementary Figure 4A-C**). Notably, two compounds known for their health-protective properties, taurine and alpha-linolenic acid (ALA), were grouped in Cluster 4, suggesting a potential link between Cluster 4 compounds and healthy aging. Due to the limited range of lipids identifiable by KEGG compounds database, Cluster 4 compounds were showing enrichment in a few terms include “glyoxylate and dicarboxylate metabolism”, “tyrosine metabolism” and others (**Figure 4D**).

**Figure 4.**
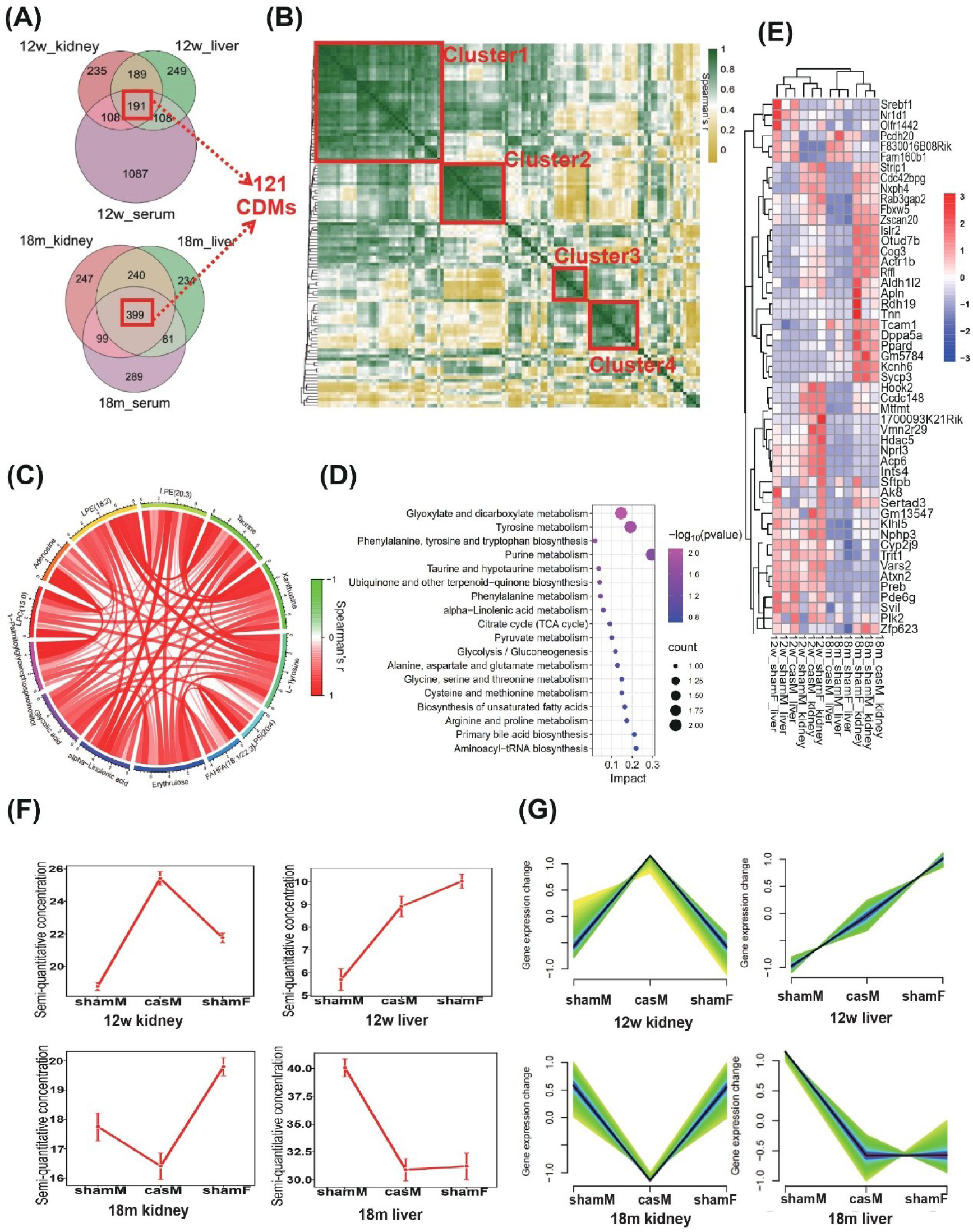
Co-expression analysis across serum, livers and kidneys identifies potential taurine-related regulators. (**A**) Venn diagram of detected compounds among serum, livers and kidneys across all the mouse groups at 12w and 18m, 121 commonly detected compounds (CDBs) were identified; (**B**) Co-expression analysis of the 121 CDBs revealed four major consensus clusters; (**C**) Correlation circos plot of the cluster 4 compounds in **B**; (**D**) Topology-based pathway enrichment analysis of the cluster 4 compounds in **B**, x-axis is the topology enrichment impact of hinted compounds; (**E**) Hierarchical cluster heatmap of z-score standardized expression levels of 51 taurine related genes in 12w and 18m kidneys and livers among shamM, shamF and casM mice. Genes selectively reduced in 18m tissues are highlighted in red; (**F**) Trends of taurine levels with standard deviation (SD) among shamM, shamF and casM across liver and kidney tissues in 12w and 18m mice; (**G**) Mfuzzing clustering of the 51 taurine co-expressed genes across liver and kidney tissues in 12w and 18m mice.

Considering the significance of taurine in aging and healthy aging, co-expression analysis and c-means clustering were applied to identify genes that positively associated with taurine. A total of 51 genes, whose expression levels showed similar dynamic trend with taurine across casM, shamM and shamF mice in kidney and liver tissues at both 12w and 18m ages, were identified (**Figure 4E-G**). Among these, 15 genes exhibited a generally decrease in expression in 18m tissues than 12w, such as *Hdac5, Preb, Plk2, Atxn2* and *Trit* (**Figure 4E**). Two key lipid metabolism genes, *Srebf1* and *Ppard* were among the 51 taurine-associated genes. *Srebf1* encodes sterol regulatory element-binding protein 1 (SREBP1), a critical transcription factor regulating lipogenes involved in fatty acid and cholesterol biosynthesis^50^. *Ppard* encodes Peroxisome proliferator-activated receptor-delta (PPAR-δ), a member of the PPAR group in the nuclear receptor superfamily, known for its ligand-binding domains (LBD) that interacts with a broad range of lipid and lipid-like molecules^51^. This includes multiple types of polyunsaturated fatty acids, such as arachidonic and linoleic acid^51^. Both PPAR-δ and SREBP1 are also involved in retinoid X receptors/ Vitamin D Receptor (RXR/VDR) pathway (**Supplementary Table 7**). Given previous research showing that intestinal-specific deletion of VDR increased taurine levels^52^, it is posited that taurine may also be associated with PPAR-δ and SREBP1 involved RXR/VDR pathway. Taken together, our results from both metabolomics and RNA-seq analysis strongly suggest that taurine is a key regulator in lipid peroxidation and metabolism, potentially playing a significant role in the process of healthy aging.

### Joint metabolomic-transcriptomic analysis reveals a stronger capacity of defensing lipid peroxidation in female and castrated male

From deep RNA-seq analysis, we identified differentially expressed genes (DEGs) among casM, shamM and shamF groups in kidney and liver tissues (**Figure 5A-B**, **Supplementary Table 6**). In the kidneys of 12w old mice, over 4500 DEGs were noted in the comparisons of "shamF *vs* shamM" and "casM *vs* shamM," while only 987 genes showed differential expression between shamF and casM. Interestingly, livers seemed to be less affected by sex and/or the changes of SH levels than kidneys as only 2053 and 2533 DEGs were detected in “shamF *vs* shamM” and “casM *vs* shamM” comparisons, respectively, in 12w-old mice, and 546 DEGs between shamF and casM. Hierarchical clustering of RNA-seq data also revealed that the transcriptomic profiles of casM samples more closely resembled those of shamF than shamM samples, with the exception of 18m-old livers, suggesting a feminizing effect of castration that creates a crossover zone between male and female transcriptomic profiles in both kidney and liver (**Figure 5C**). However, the impact of sex and/or SH levels on transcriptomes, pronounced at 4 weeks post-castration, appeared to diminish in 18-month-old mice. In this older age group, the numbers of DEGs among the 3 groups were reduced, especially the comparisons of shamM with casM. In 18-month-old kidneys and livers of “casM *vs* shamM”, it showed only 435 and 353 DEGs, respectively.

**Figure 5.**
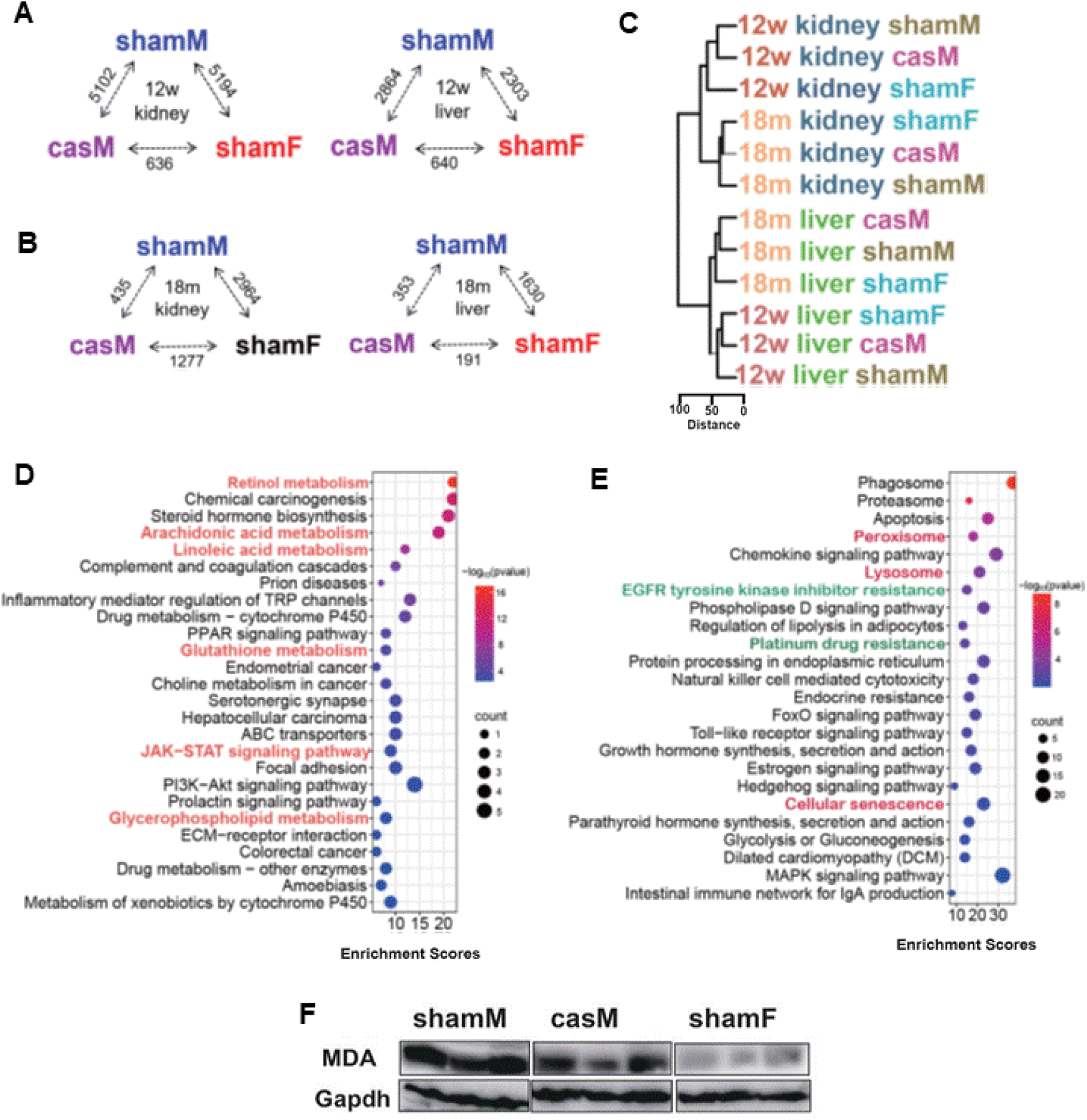
Joint metabolomic-transcriptomic analysis reveals a stronger capacity of defensing lipid peroxidation in female and castrated male. (**A-B**) The numbers of DEGs among livers and kidneys tissues of shamM, shamF and casM mice collected at 12w (**A**) and 18m (**B**); (**C**) Hierarchical clustering of 12w and 18m liver and kidney samples demonstrating casM’s closer alignment with shamF than sham; (**D, E**) Representative significantly enriched KEGG pathways in 18m liver samples from ’casM *vs* ShamM,’ (**D**) and ‘shamF *vs* ShamM’ (**E**), identified by joint transcriptomic-metabolomic analysis using DEGs and SAMs. Pathways marked in orange indicate common terms between these two comparisons. (**F**) Immunoblotting of MDA in 18m livers;

Mapping the DEGs onto mouse chromosomes, we found that DEGs in "shamF *vs* shamM" and "casM *vs* shamM" from both 12w and 18m-old liver tissues were significantly clustered to chr4B3 (FDR < 0.05). Of the 145 annotated genes in chr4B3, 46 belong to the Major Urinary Proteins (*Mup*) family, with 25 being pseudogenes. Except for *Mup4* and *Mup5*, which were barely detectable, the other *Mup* family genes in chr4B3 were drastically reduced in casM and shamF livers, showing an average log2FC = -5.57 (**Supplementary Figure 5A**). *Mup14*, *7*, *11*, and *1* were the top 4 differentially expressed *Mup* genes. It is known that *Mup* genes, predominantly expressed in the liver, exhibit sexually dimorphic traits. Male mice typically excrete 2– 8 times more Mup protein in their urine compared to females ^53^. Our results indicate that the expression of *Mup* genes is influenced not only by sex genotypes but also by testosterone levels.

To identify potential long-term salutary regulatory elements brought by sex and/or castration at old age, we conducted a joint metabolomic-transcriptomic analysis. Here, shamM mice, typically having a shorter liferspan, served as controls, while shamF and casM mice, associated with healthier and longer lifespans, were the focus ^54^. We utilized the SAMs and DEGs from “shamF *vs* shamM” for the joint analysis to investigate female-specific signatures; meanwhile, the joint metabolomic-transcriptomic analysis was also performed using the SAMs and DEGs from “casM *vs* shamM” for interpreting castration-specific features. In 18m-old kidneys, both shamF and casM groups showed enrichment in KEGG programs related to “lysosome” and “cholesterol metabolism” (**Supplementary Table 8-9**). In the liver of the same age group, the significantly enriched KEGG pathways in the “casM *vs* shamM” comparison are illustrated in **Figure 5D**. Notably, 20 terms, such as “linoleic acid metabolism”, “arachidonic acid metabolism”, “retinol metabolism”, “Jak-stat3 signaling” and “glutathione metabolism” were consistent with those in the “shamF *vs* shamM” comparison (terms marked by orange in **Figure 5D and 5E**, **Supplementary Table 10-11**). These KEGG terms are known to play critical roles in lipid peroxidation.

To further validate these findings, we measured levels of malondialdehyde (MDA) -a final product of polyunsaturated fatty acids peroxidation, in the three groups of mice. Results indicated that shamF mice exhibited the lowest, while shamM mice had the highest MDA levels in 18m-old livers, suggesting that females and castrated males experience less lipid oxidation stress than their male counterparts (**Figure 5F**). Additional oxidative stress markers, SOD1 and SOD2, were also tested and showed similar trends to MDA among three groups (**Supplementary Figure 5B**). Taken together, our analysis suggests a close association between lipid metabolism and sex and SH levels. CasM and shamF mice are likely to absorb more health-beneficial compounds from their diet and experience reduced lipid peroxidative stress, potentially contributing to their longer lifespans.

### Evaluating sex-dependent cisplatin toxicity through castration models

The clinical application of anti-tumor drugs, particularly Platinum-based drugs (PBDs) like cisplatin, which are used in nearly half of all chemotherapy treatments^55^, is often hindered by drug-induced tissue toxicity. Notable adverse effects of these drugs include nephrotoxicity and hepatotoxicity, exhibiting differential impacts between male and female patients^56^. Such sex-dependent disparities in toxicity may be influenced by SHs, as suggested by the increased risk of nephrotoxicity observed in perimenopausal women undergoing cisplatin treatment^57^. Aging, a key factor influencing SH levels in both sexes, further complicates this relationship. To evaluate the utility of the castration model in studying the role of SHs in sex-dependent toxicity, we treated 12w-old mice (4 weeks post-castration/sham procedure) with a low dose of cisplatin twice, every 72 hours. Given that the kidney, and to a lesser extent the liver, are particularly vulnerable to cisplatin-induced damage, we conducted histological analyses on the renal cortex and left liver lobe tissues post-treatment.

Hematoxylin and eosin (H&E) staining of liver tissues showed the most severe liver injury in shamF mice, characterized by marked steatosis, sinusoidal congestion, and hemorrhage. In contrast, sham-operated male (shamM) and castrated male (casM) mice displayed comparatively less hepatic damage (**Figure 6A**). Conversely, Periodic Acid Schiff (PAS) staining of kidney tissues showed that casM mice had more severe kidney damages than both shamF and shamM, with a notable increase in acute renal tubule injury (**Figure 6B**). Previous studies on cisplatin-induced hepatotoxicity have yielded limited insights regarding sex differences. However, substantial evidence indicates that males are more susceptible to cisplatin-induced nephrotoxicity^56^. Studies administering cisplatin to rats at both 2 weeks (1 mg/kg/day) and a single dose (7.5 mg/kg) have shown that males experience more severe nephrotoxic effects than females^58^. Our pathological observations, combined with previous findings and the fact that castrated male (casM) mice have the lowest testosterone (TS) levels, suggest that intermediate TS levels, similar to those in shamF mice, may offer greater protection against renal toxicity induced by cisplatin.

**Figure 6.**
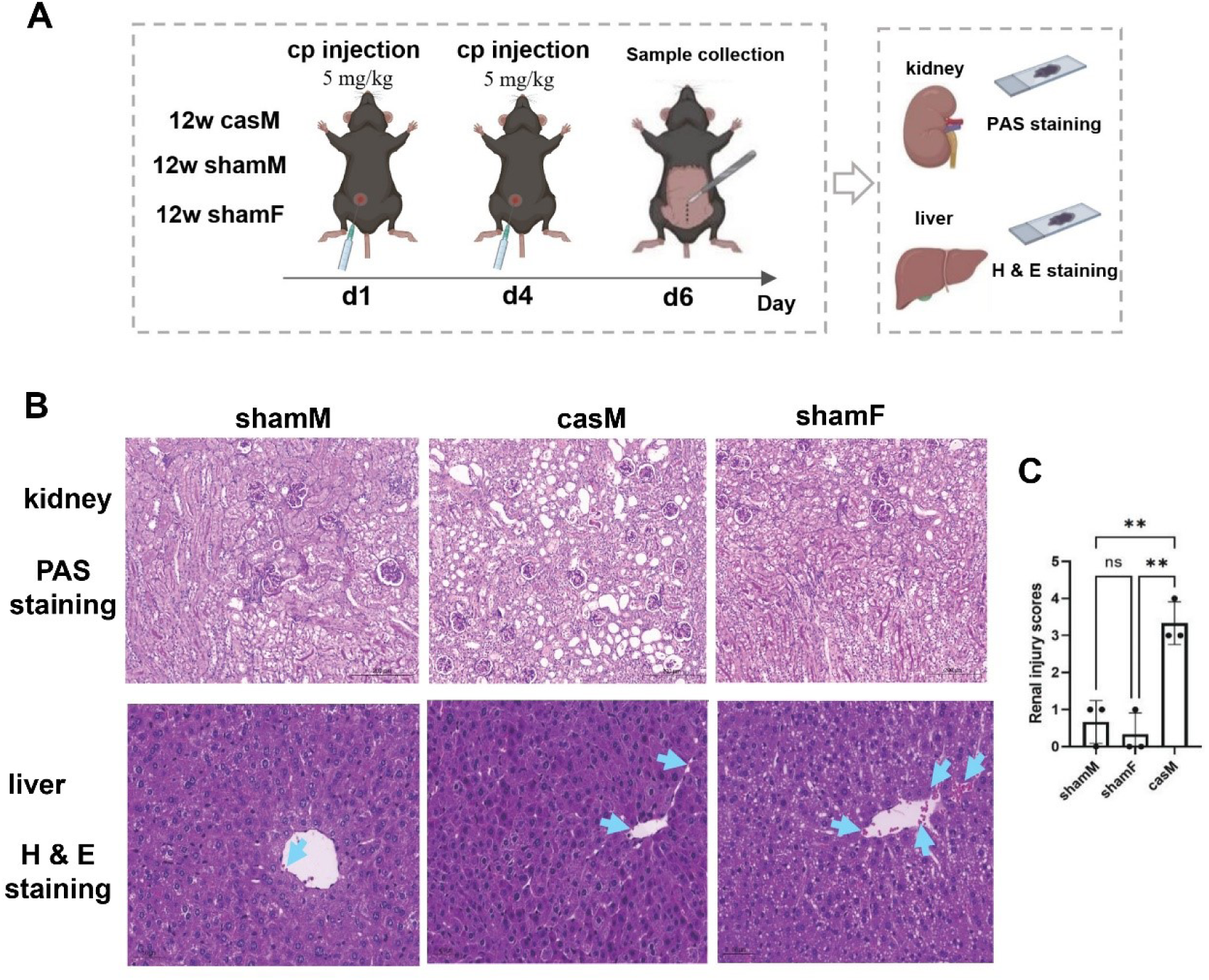
Illuminating sex hormone influence in cisplatin toxicity through castration models. **(A)** Graphic illustration of the experimental design; (**B**) PAS staining of kidneys (upper panel) and H & E staining of livers (lower panel) in 12w mice (4 week post castration/sham procedure) with cisplatin treatment, blue arrows indicate representative injury spots in livers; (**C**) Bar plots of renal injury scores in three groups of mice by One-Way ANOVA test (Tukey correction was used for correction of multiple comparisons, \*\**p* < 0.01, ns: not significant).

### Evaluating castration model in sex-dependent cognitive function

Increasing evidence has revealed that cognitive function decline is sex-dependent, for instance, a higher frequency of Alzheimer’s diseases (AD) is reported for women than men: nearly two-thirds of patients with AD are women^59^. However, the roles of SHs in neurodegeneration process is rarely explored. Given the known correlation between ages 10 – 14 months in mice and 38 – 47 years in humans, where senescence-related changes become prominent^60^, we conducted aging-related behavioral analyses among the three groups of mice, to detect the potential roles of castration in cognitive function. Barnes maze (BM), open field (OF) and novel object recognition (NOR) tasks were tested to evaluate the spatial learning and memory, locomotor activity, and recognition memory in 12 – 13 month-old casM, shamM and shamF mice (**Figure 7A**). Previous studies have shown pronounced cognitive decline in middle-aged mice (8 – 12 months) as assessed by the BM test^61^. However, our results showed that middle-aged casM demonstrated superior performance, spending less time and making fewer errors with less immobile time in locating the escape hole than shamM and/or shamF (**Figure 7B**). Non-significant difference was found in OF and NOR tasks across the three groups (**Supplementary Figure 6A-B**). Our results revealed that deprivation of TSs by castration procedure can confer a protective effect against cognitive decline. This selective enhancement in spatial learning and memory may reflect the differential influence of castration on brain regions, particularly affecting areas like the hippocampus that are crucial for navigation tasks, while exerting less impact on regions involved in locomotor and recognition memory functions.

**Figure 7.**
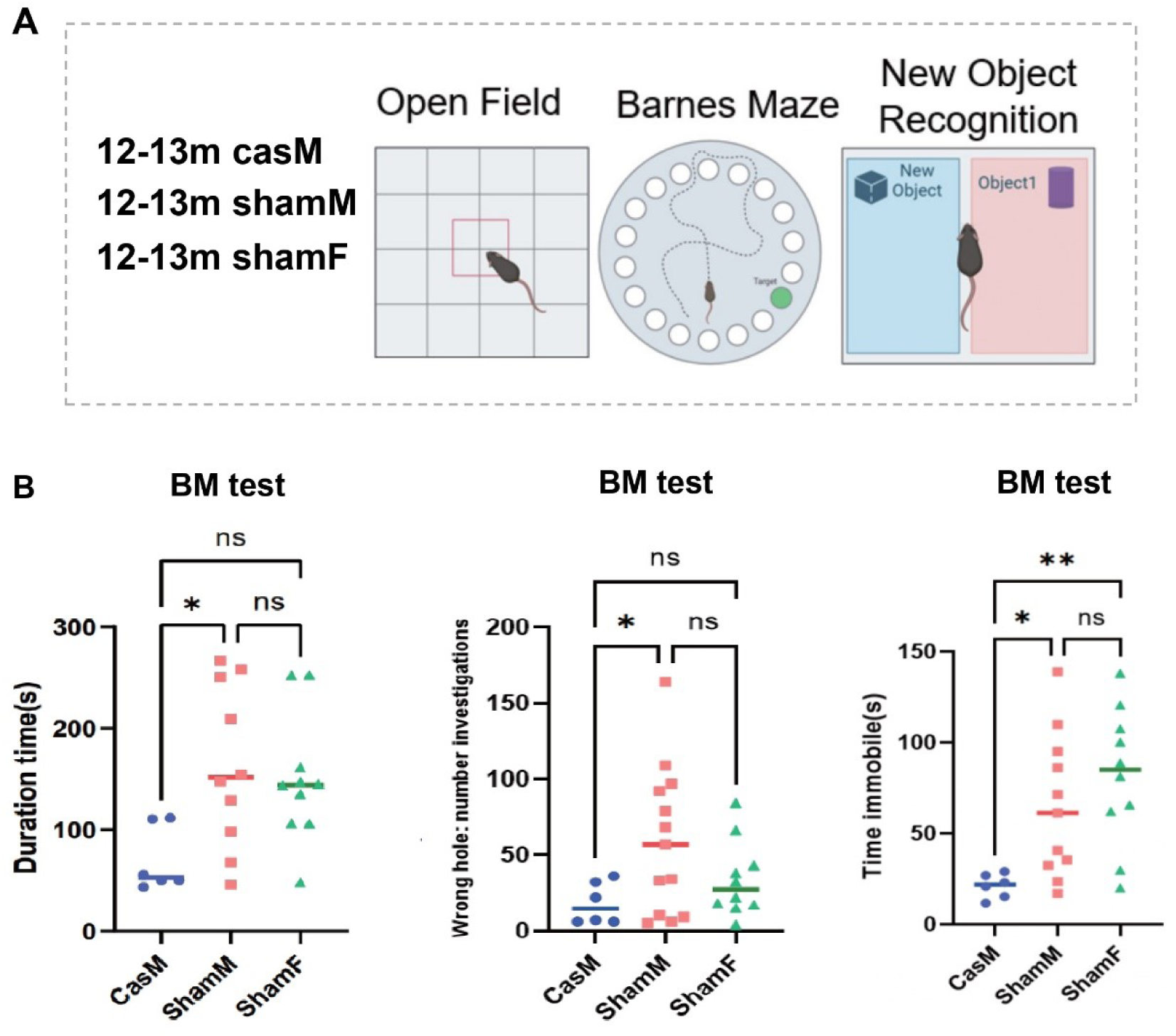
Aging-related cognitive function tests among three groups of mice. **(A)** Graphic illustration of the behavioral tests; (**B**) Analysis of BM tests results: duration time that mice have spent to find the correct target escape hole (left panel), exploration of wrong holes numbers (middle panel), and immobile time during the test (right panel) among casM (n=6), shamM (n= 13) and shamF (n = 10) by One-Way ANOVA, *\*p* < 0.05, *\*\*p* < 0.01, ns: not significant. For Figure 1C & E-G, Tukey correction was used for correction of multiple comparisons and median lines were shown.

## DISCUSSION

Recognizing and addressing sexual differences in both basic and clinical research is critical, yet historically overlooked. On one hand, the bias toward using male subjects in biomedical research has often been justified by the desire to minimize variability, particularly traits influenced by the female estrous cycle^62^. This male-centric focus has inadvertently limited our understanding of tissue function and disease mechanisms from a sex-specific perspective, thereby highlighting the pressing need for more inclusive animal models. On the other hand, straightforwardly comparing male with female is generally applied in the increasingly emphatic but still limited sex-difference studies, which made the leading mechanisms hard to recognize due to complicated variables between male and female in both genetic and non-genetic aspects.

C57BL/6 mouse strain is one of the most widely used models in biomedical research ^63^. A systematic exploration of sex-and hormone-related molecular features in this strain is crucial for advancing our understanding of sex-biased biological processes. Moreover, it is critical to evaluate the utility of the castrated C57BL/6 model in addressing sex-specific research questions.. In this study, we integrated observational investigations, behavioral tests and multi-omics approaches to systematically investigate the differences among shamM, casM and shamF C57BL/6 mice across multiple tissues at different stages. We found that adult casM mice spontaneously consumed less food than shamM, a behavior we believe to be the primary factor contributing to their reduced rate of body weight gain. Considering the well-established connection between dietary restriction, body weight, overall health, and longevity^64, 65^, it is plausible to suggest that this dietary pattern confers benefits for healthy aging in casM mice. The underlying triggers prompting reduced food intake in casM mice merit further investigation. Intriguingly, despite identical diet, shamF and casM mice exhibited elevated levels of multiple food-derived health-promoting compounds compared to shamM, including taurine, a compound that recently known for its aging-slowing effects in various species^36^. These findings point to a potential influence of the gut microbiome in modulating host metabolism and appetite regulation^66^, which in turn can affect eating behaviors and the bioavailability of beneficial compounds in food. Given the dynamic interplay between the gut microbiome, aging, and its sex-dependent modulation^67, 68^, our results hint at the involvement of sex hormones in shaping the microbial community. This, in turn, could influence dietary behaviors and the assimilation of protective compounds from food. Future research delving into the microbiota-gut-brain axis and its relationship with sex difference and healthy aging in this castration model is warranted.

To broaden the feasibility and practicability of applying the castration model in elucidating the molecular mechanisms of sex differences, we investigated the differences in tissue injuries induced by tumor drugs and the cognitive differences among three groups of mice. While more severe renal damages have previously been observed in males compared to females following cisplatin treatment at both young and old ages^69^, the specific roles of genetic and non-genetic co-determinants in this disparity are still elusive. Rostami et al. reported that rats, 1 week post-castration and treated with a low dosage of TS rather than high dosage, exhibited increased tolerance to cisplatin nephrotoxicity^70^. Complementing this, our pathological assays showed that castration alone, without TS treatment, exacerbated renal toxicity. These findings collectively suggest that balanced levels of TS are essential in defending against such toxicity. Given that aging is a major contributor to changes in SH levels in both sexes^71, 72^, and considering the significant impact of these changes on chemotherapy-induced toxicities, there is an imperative need for more comprehensive research to integrate these hormonal alterations with chemotherapy toxicity across various age groups, thereby enhancing the understanding and management of chemotherapy side effects in the aging population.

Previous research has offered mixed insights into the impact of castration on cognitive function. One such study employed a dual-task approach – utilizing an 8-arm radial maze and a step-down avoidance task – to assess spatial and short-term memory in rats. Conducted approximate 82 days post-surgical and chemical castration, the study concluded that castration deteriorates memory function^73^. In contrast, our findings, based on BM test in castrated mice, suggest an enhancement in cognitive performance. Several factors could account for these divergent outcomes. First, the timing of cognitive assessments post-castration differed between the studies. Our behavioral tests were performed on 12-13month old mice, nearly 10-11 months after castration or sham operations. It is plausible that the benefits of castration on spatial memory may not be readily apparent in younger rodents but become more pronounced with age. Second, the choice of behavioral tasks could also contribute to the observed discrepancies. The step-down inhibitory avoidance task, employed in the previous study, uses immediate electric shocks as stimuli to evaluate aversive memory, while the 8-arm radial maze task requires deprivation of food and/or water for animals. Electric shock is a type of psychological stress for rodents^74^. Mounting evidence suggests that testosterone significantly modulates physiological responses to stressors^75^; thus, the markedly reduced testosterone levels in castrated mice could potentially confound the evaluation of cognitive performance. To mitigate potential confounders, we opted for less stressful tasks – namely BM, OF and NOR tests. Our results from the BM test indicate that castration may serve as a good model to study the roles of sex hormones in the sex difference of cognitive function.

In summary, our study underscore the versatility of castration mouse model in sex-dependent biological queries. The model effectively "feminizes" males at multiple molecular levels through alterations in SHs, while preserving the somatic male genotype. This unique duality positions the castration model as a useful tool for distinguishing between hormone-induced sex differences and genotypic sex, broadening its utility beyond the scope of sex-biased research.

## METHODS

### Animals, surgical castration procedures, and sample collections

C57BL/6 mice were purchased from Zhuhai BesTest Bio-Tech Co,.Ltd. and housed in the animal facility of Shenzhen TopBiotech Co., Ltd (Shenzhen, China) under pathogen-free condition with free access to chow and water. Male and female mice at 8 weeks old were anesthetized by isoflurane for castration (testicles removed) or sham procedures. Mice were placed on a warm pad after suture, then transferred to cages while awake. 4-weeks post-castration/sham operations, serum was collected for steroid hormone detection and untargeted metabolites analysis while liver and kidney were dissected for untargeted metabolites analysis and RNA-seq (n=3-6/group). Castrated/sham mice at 12-13m old were subjected to behavioral tests and were sacrificed for sample collection at 18m. At 18m, serum was used for untargeted metabolomics while liver and kidneys were dissected for both untargeted metabolomics and RNA-seq (n=3-6/group). The animal study protocol was approved by the Animal Ethics Review Board of Shenzhen TopBiotech Animal Facilities (No. TOP-IACUC-2021-0100).

### Food intake, body length and rectal temperature measurement

To investigate the potential link between reduced food intake and slower body weight gain in castrated male mice (casM), we monitored the weekly changes in chow weight over a 4-week period in 3 cages per group, each housing 3-5 mice aged 27 weeks. The average food intake per mouse was calculated on a weekly basis. In assessing the impact of castration on growth, we measured the body lengths of 29-week-old casM (n = 13), shamM (n = 12), and shamF (n = 13) mice. Mice were gently extended to full length, and measurements were taken from apex nasi to the base of the tail (tail not included). Additionally, rectal temperature was measured on 30 weeks and 32 weeks old casM (n = 13), shamM (n =12) and shamF (n = 13) mice, with the data combined for subsequent analysis. Statistical differences in food intake, body length, and rectal temperature across the three groups were evaluated using one-way ANOVA, multiple comparisons with Tukey correction.

### Sample treatment for targeted steroid hormone spectrum detection

50 μL serum was thawed and vortexed for 30s following by mixture with purified H_2_O, internal standard working solution and methanol. The well mixed samples were then subjected to repeated 13000 rpm centrifuge for 10 minutes (mins) under 4°C. The supernatant was retained further purified with OASIS PRiME HLB uElution (SPE). 200 μL methanol and 200 μL H_2_O were separately used to wash and equilibrate the SPE cartridges. The supernatant was loaded to the equilibrated cartridges and then washed by 200 μL 10% ACN/H_2_O (v/v) and 200 μL Hexane. The flow-through fraction was discarded. Rinse the cartridge was with 40 μL 90% ACN/H_2_O (v/v) and collect the cluent. Add 60 μL H_2_O to the cluent and shake the mixture for 3 mins. The final mixture and the same volume of isotopically-labelled internal standards were subjected to UHPLC-MS/MS analysis.

### Serum sample preparation for untargeted MS analysis

150 μL methanol which was premixed with isotopically-labelled internal standard compounds was added to 50 μL mouse serum and vortexed for 30 seconds to mix well. The mixture was then sonicated for 10 mins in ice water bath and incubated for 1 h at -40 ℃ to precipitate proteins. After 15-min high-speed centrifugation, the supernatant was used for MS analysis. An equal aliquot of the supernatants obtained from each serum sample was mixed as the quality control (QC).

### Kidney/liver sample preparation for untargeted MS analysis

10-25 mg tissues were submerged with extraction solution (methanol:water = 3: 1, with isotopically-labelled internal standard mixture) and homogenized in ice-water bath. Incubate homogenized samples for 1 h at -40°C and centrifuge at 12000 rpm for 15 min at 4°C. The supernatant was retained for MS analysis. A volume matched quality control (QC) sample was prepared for MS.

### UHPLC-MS/MS analysis for targeted steroid hormone spectrum

EXIONLC System (Sciex) equipped with a Waters Acquity Uplc Bch C8 column was utilized for UHPLC analysis at Biotree Technology Co., Ltd, Shanghai, China. Assay development was applied by SCIEX 6500+ triple quadrupole mass spectrometer (Sciex) with an IonDrive Turbo V electrospray ionization (ESI) interface, under multiple reaction monitoring (MRM) module. MassLynx Work Station Software (Version 4.1) was employed for MRM data acquisition and preprocessing.

### UHPLC-MS/MS untargeted metabolomics analysis

T3 chromatographic column with affinities to both lipid and hydrophile at Biotree Technology Co., Ltd (Shanghai, China) was used to acquire the metabolome of castrated male (casM), sham male (shamM) and sham female (shamF) mice by Orbitrap mass spectrometer. Joint negative (NEG) and positive (POS) ionization modes were employed for detecting metabolites based on their mass to charge ratios. Raw data was converted to the mzXML format using ProteoWizard platform (https://proteowizard.sourceforge.io/) and processed with an in-house R script and XCMS tool^76^ for peak detection, extraction, alignment, and integration. Data was subjected to fragment extracted ion chromatogram (MS^2^) annotations as previously described^77^, MS^2^ metabolites with scores > 0.3 were retained for next-step analysis. These compounds were mapped to Human Metabolome Database (HMDB) and classified into multiple categories. The biggest category under both NEG and POS modes is lipid and lipid-like molecules, which counts for ≥ 30% identified metabolites (**Figure 2A-B**, **Supplementary Table 2-3**). Compounds from NEG and POS modes were merged for down-stream analysis.

### RNA extraction for RNA-seq analysis

Total RNA was extracted using Trizol reagents (Invitrogen Cat#15596026) as we have described before^78^. PolyA-tailed RNA was enriched for constructing 150bp pair-end RNA-sequencing libraries using MGISEQ-2000RS kit (MGI Cat#1000012555), and sequenced by DNBseq MGI2000 platform at BGI genomics Co., Ltd, Shenzhen, China.

### Cisplatin administration

Mice were administrated for intraperitoneal injection of cisplatin (Cat# 232120, sigma) at 4 weeks post-castration/sham operations, with a dosage of 5 mg/kg every 72 hours for 2 times. Mice were sacrificed 48 hours after cisplatin treatment for the collection of liver and kidney tissues.

### Histopathological analysis

For liver and kidney tissue examination, samples were first fixed in 10% formalin solution for 48 hours, followed by paraffin embedding. Sections of 3 μm thickness were then prepared for histological analysis. Hematoxylin and Eosin (H & E) and Periodic Acid-Schiff (PAS) staining were performed according to established protocols. Liver injury was evaluated by visually inspecting the patterns of steatosis, sinusoidal congestion, and hemorrhage. Renal injury was quantitatively assessed using the scoring criteria established in a previous study^79^. Statistical differences of renal injury scores across the three groups were assessed one-way ANOVA (multiple comparisons with Turkey correction).

### Western-blot

Total protein was isolated on ice by cell & tissue lysis buffer (Cat#P0013, Beyontime, China) and quantitated using BCA assay kit (Cat#P0012, Beyontime, China). Approximate 15ug protein was used for western-blot assays following previous protocols^80^. Antibodies for MDA, SOD1 and SOD2were purchased from abcam (Cat# ab243066, ab13498 and ab13533), GAPDH was purchased from Beyontime (Cat# AF0006).

### Behavioral tests

Three groups of mice include casM, shamM and shamF at 12-13 months old (n= 6-13) were subjected to BM, OF and NOR tasks. A 60 cm high, 120 cm-diameter and 60 mm thick circular wheeled PVC slab with 25 cm-diameter holes was used for BM tests. A modified protocol was employed following the standard methods in previous study^81^. Briefly, mice were firstly trained in continuous 7 days to find the escape hole within 4 mins; if failed, mice were finally guided to the target hole gently. Mice were subjected to the probe trial on day 8. OF test was applied to test the exploratory behavior and general activity of mice, mice were placed in an enclosure, rectangular tank with surrounding walls for 5 mins, distance moved, time spent moving, rearing, and change in activity over time for each mouse were recorded. NOR test was performed following previous study^82^. Mice were placed in the empty tank for 10 mins on day1; day 2, two objects were placed in the tank, and mice were allowed for 10 mins to explore; one of the objects was replaced by a new one on day 3, and time spent for exploration (TE) within 5 mins for each object was recorded. NOR index was calculated via this formulation: NOR index = new object TE-old object TE/ Total TE. Mice behaviors during these tests were recorded and analyzed by ANY-MAZE software (https://www.any-maze.com/). Statistical differences in BM, OF and NOR tasks among the three groups were analyzed via one-way ANOVA, multiple comparisons with Turkey correction.

### Identification of significant altered metabolites (SAMs) and differentially expressed genes (DEGs)

A strict pipeline was applied to identify SAMs. Firstly, the normalized peak values of metabolites were subjected to unsupervised principal component analysis (PCA) and then administrated to supervised orthogonal projections to latent structures-discriminate analysis (OPLS-DA) for data modeling by commercial SIMCA software (V16.0.2). A 7-fold cross validation was employed for model selection. To evaluate the predictive power and robustness of OPLS-DA model, a 200-times-permutations was performed and the value of variable importance in the projection (VIP) of the first principal component in OPLS-DA analysis was calculated which represents the contribution of each variable to the model. Metabolites with VIP > 1 and students’ t-test *p*-value < 0.05 were recognized as SAMs. For DEG identification, quality control was performed for raw RNA-seq reads by SolexaQA (-h 30, -l 30)^83^. Cleaned reads were subjected to Hisat2^84^ for mapping. Cufflinks package^85^ was utilized to calculate the Fragments Per Kilobase of transcript per Million mapped reads (FPKM) values and to identify DEGs use FDR < 0.05 as the cutoff value.

### Expression trend clustering analysis

To identify expression trend patterns of metabolites among the serum samples of 12w casM, shamM and shamF, the union of SAMs across three pairwise comparisons in **Figure 3E** were analyzed by Mfuzz package (c=9, m =1.25)^86^. To illustrate genes that potentially related with taurine, the expression levels (FPKM values) of all the annotated genes in kidneys and livers at 12w and 18m casM, shamM and shamF mice, were also subjected to Mfuzz package^86^ for clustering (c=9, m =1.25). The semi-quantitative concentrations of taurine were plotted as stack lines. Genes that show similar expression trend with taurine in all the tissues were retained (**Figure 4F-G**).

### Hierarchical clustering analysis

Hierarchical clustering analysis for SH levels after castration/sham treatments (Figure 2A), the expression of 51 taurine related genes (**Figure 4E**), serum metabolomic profiles among casM, shamM and shamF using (**Figure 3B**) and tissue RNA-seq gene expression profiles (Figure 6C), were performed by Hclust function in R platform using Euclidean distance metric and complete lineage. The z-score standardized SH levels and expression levels of taurine related genes after clustering were visualized via heatmap using pheatmap 1.0.12 package.

### Enrichment analysis of SAMs

SAMs were subjected for topology-based KEGG pathway enrichment analysis using MetaboAnalyst with default parameters^87^. Differential abundance (DA) scores^88^ were calculated for the significant enrichment terms using this formula: DA scores = 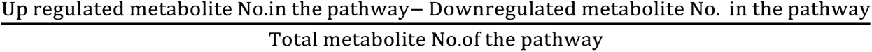

### Joint metabolome & transcriptome analysis

To examine the interplay between metabolomics and other transcriptomics layers, SAMs and DEGs were used for joint metabolome & transcriptome enrichment analysis against KEGG pathways via MetaboAnalystR under joint-analysis module by hypergeometric test with default parameters. FDR < 0.05 was applied as the cut-off value

## Author Contributions

J.J. conceived and designed this study; J.J. and W.W. wrote and revised this manuscript; S.Y. and F.L. provided necessary resources to this study; J.J. performed the metabolomics, RNA-seq analysis, and the western blot assay; N.G. and X.G performed the metabolomics analysis; Y.W. and G.W. performed the animal behavior tests; J. Q., and K. G. performed the sham/castration procedure; J.J., Y.W., G.W., J. Q., K. G., X.G and Y.L. did the mouse colony and tissue samples collection.

### Acknowledgement

We acknowledge Ms. Jiahui Liu for her generous help for sample collection and Dr. Xinhui Liu for his kind help for assessing renal injury scores. This work was supported by Shenzhen Science and Technology Program (Grant No.RCBS20210706092341003) to J.J.

