## Supplementary Figures & Tables for "Applicability of Castration Model in Sex Difference Studies: Insights from Metabolome and Transcriptome Analyses": 20240904_Supplementary Information.docx

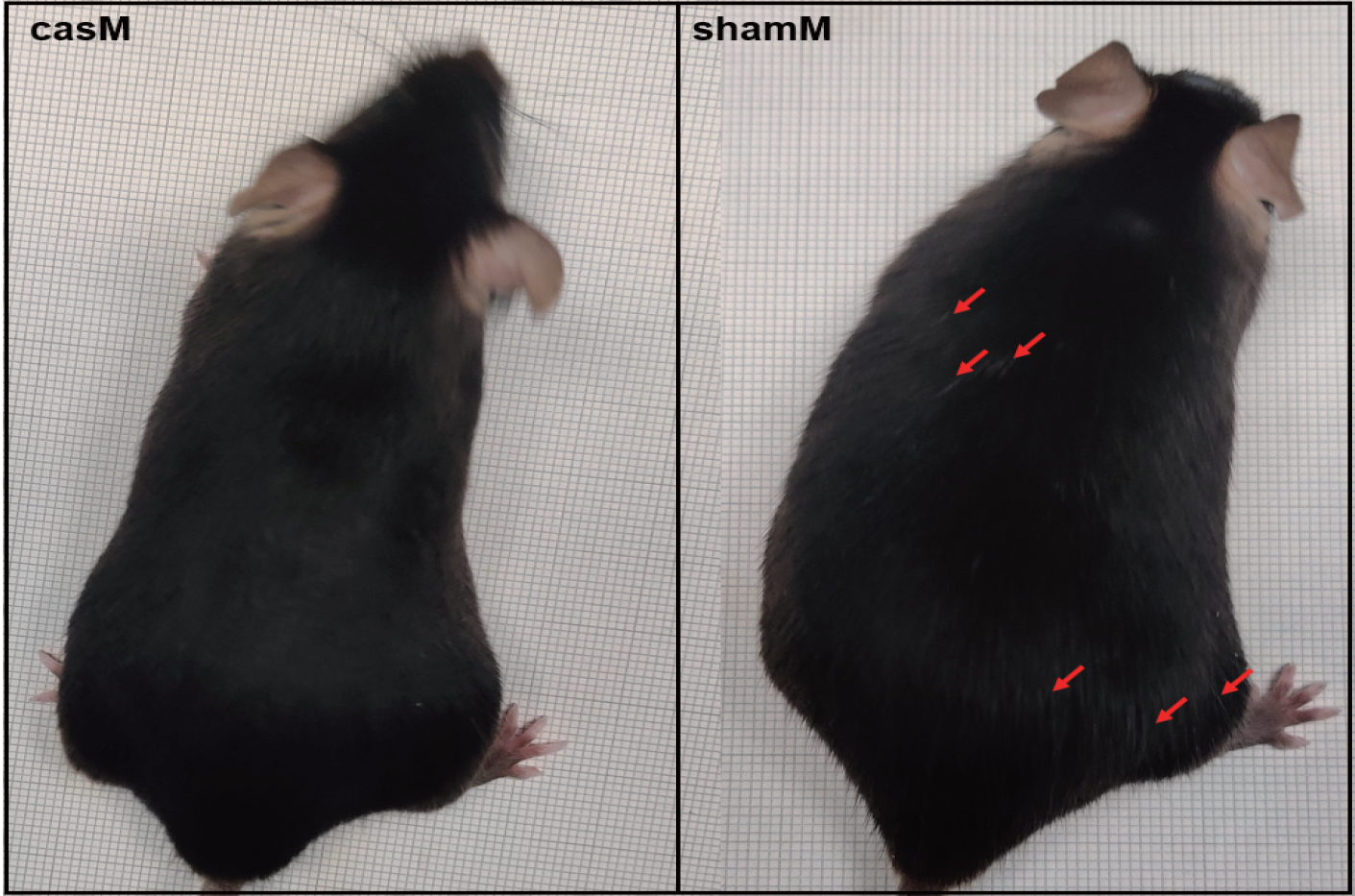

**Supplementary Figure 1** Castration or sham procedure was processed in shamM than casM at 8 weeks old, and more gray hairs (see red arrows) were observed in shamM than casM at 12-13 months old**.**

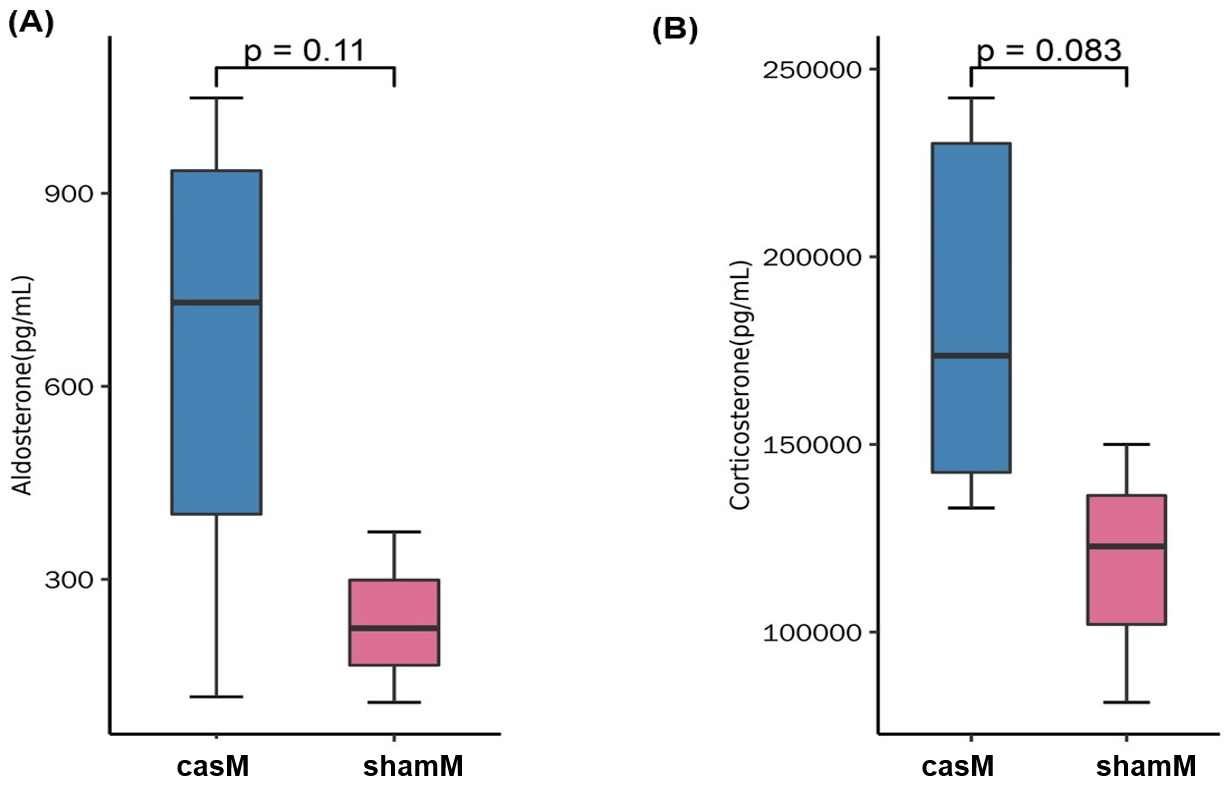

**Supplementary Figure 2** Slight increase of aldosterone (A) and corticosterone (B) was observed in casM than shamM (unpaired students’ t-test, *p*-values were shown on top).

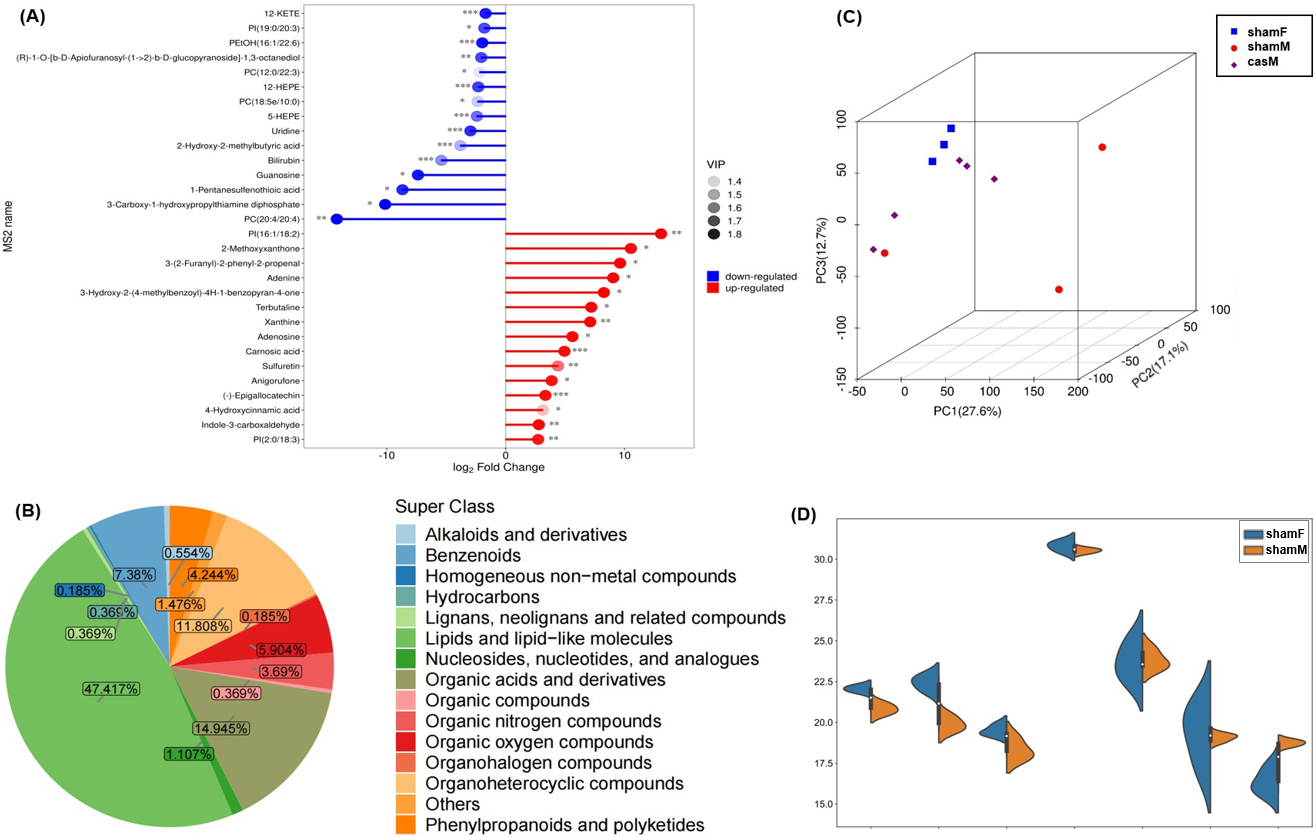

**Supplementary Figure 3 (A**) Matchstick plot of the top SAMs between 12w shamM and shamF (ranked by fold changes, **p* < 0.05, ***p* < 0.01, ****p* < 0.001); (**B**) Compounds categories of 18m mouse serum; (**C**) 3D PCA analysis of casM, shamM and shamF using the 18m serum metabolomics profiles; (**D**) Comparisons of some selected compounds between 18m shamM and shamF, from left to right are D-pantothenic acid, ruscogenin, 2-hydroxycinnamic acid, arachidonic acid, 7-ketocholesterol, 12-HEPE and 5-HEPE.

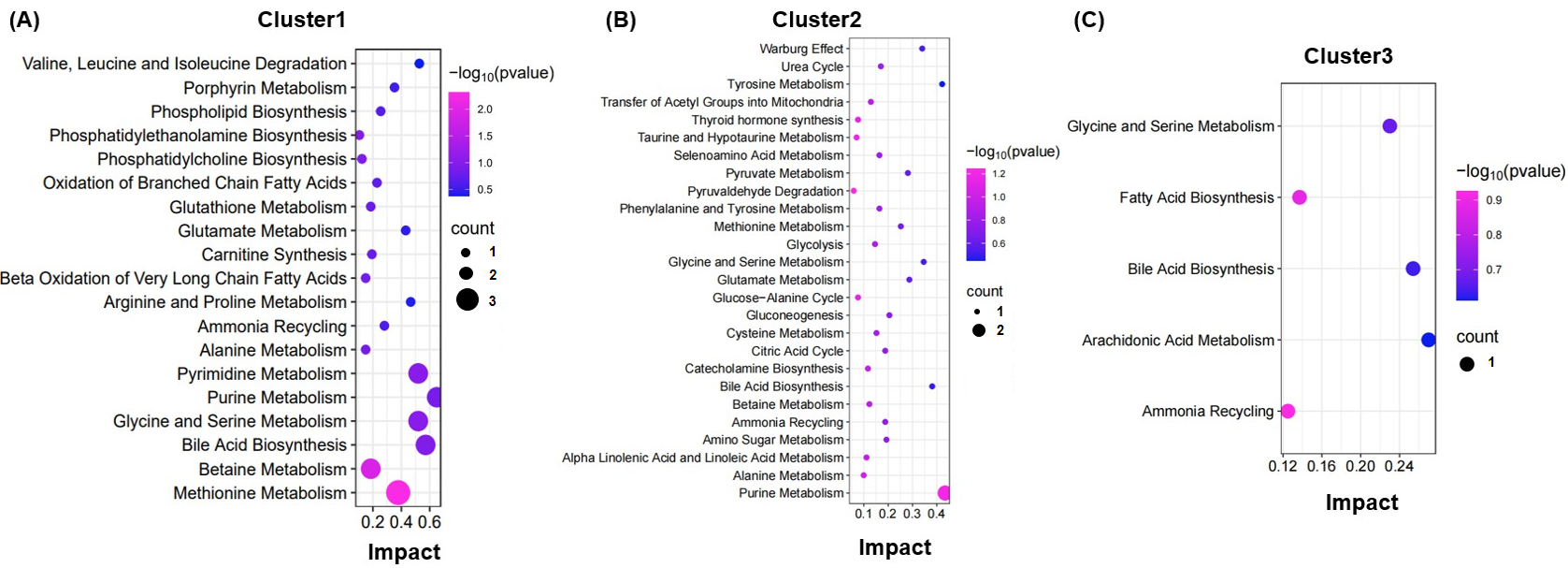

**Supplementary Figure 4. (A) - (C)** Topology based enrichment analysis of consensus clusters 1-3 identified in the co-expression analysis of 121 CDBs in Figure 4B**.**

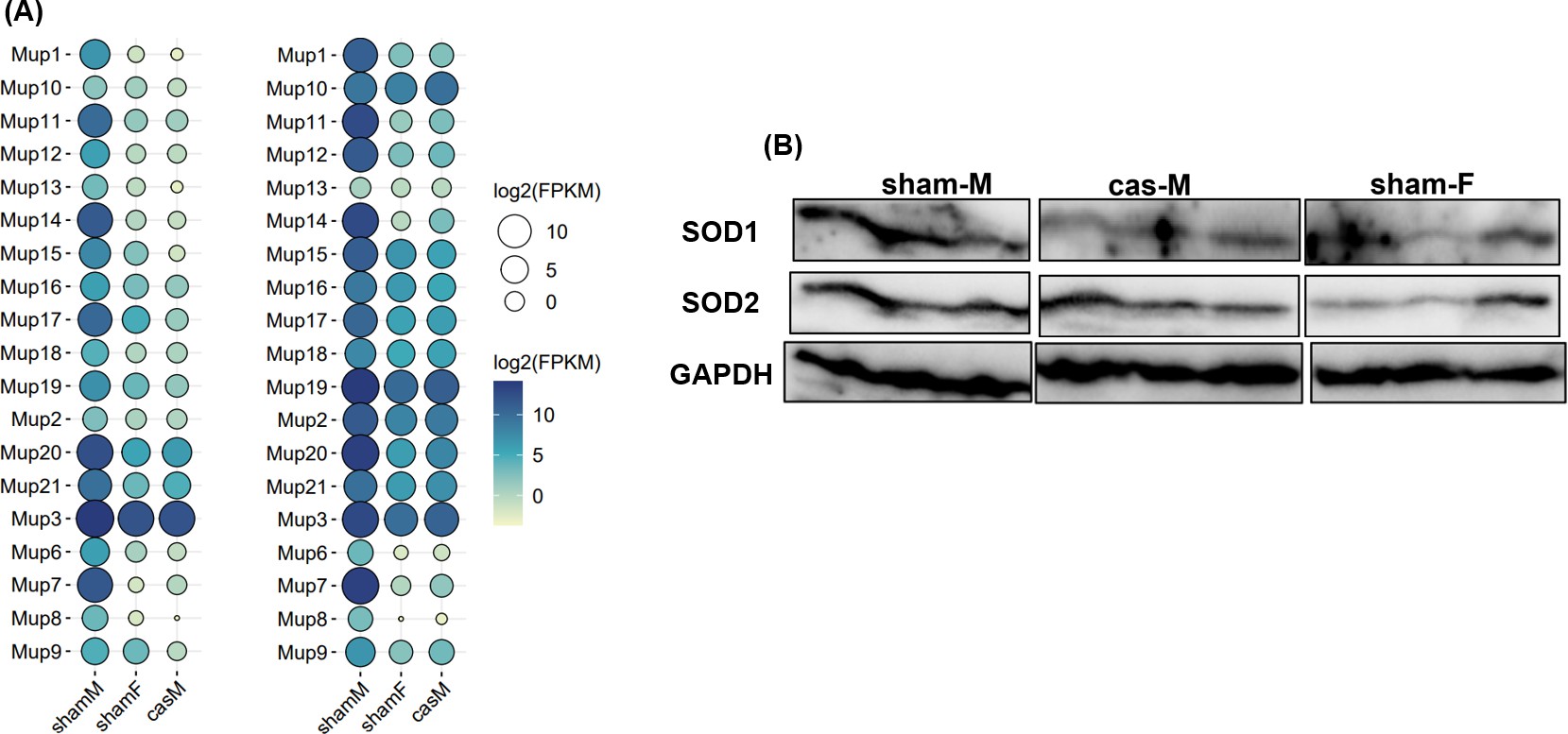

**Supplementary Figure 5. (A)** Bubble plot of *Mup* gene family expression of liver tissues among shamM, shamF and casM mice. Left, 12w-old liver, Right, 12m-old liver. **(B)** Western blot of SOD1 and SOD2 among the liver tissues of 18m shamM, shamF and casM mice.

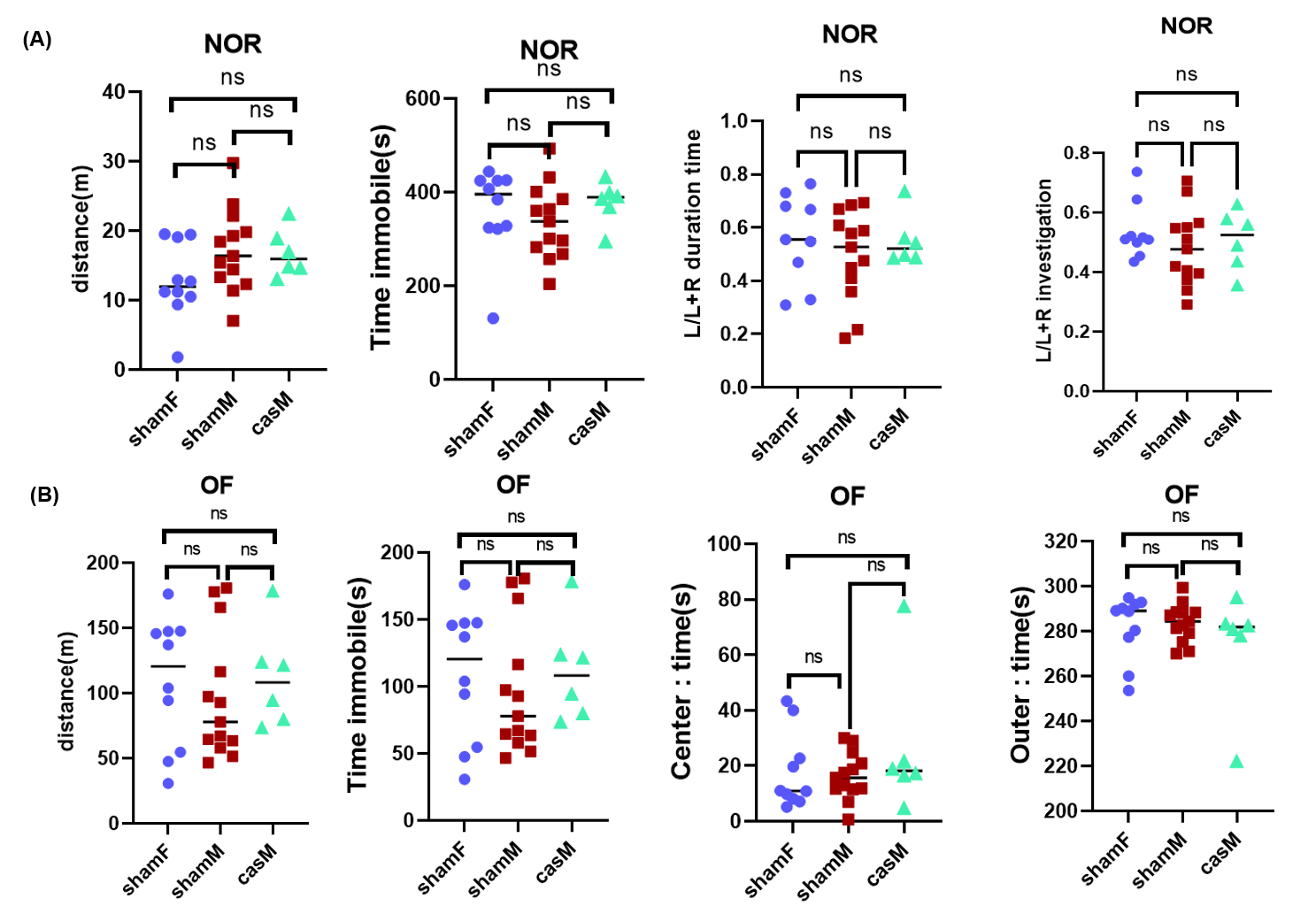

**Supplementary Figure 6.** ShamF, shamM and casM mice were subjected to sham or castration procedure at 8 weeks old and these 3 groups of mice show non-significant difference in (**A**) new object recognition (NOR) and (**B**) open field (OF) behavioral tests at their 18 months old (One-Way ANOVA analysis, Tukey correction was used for correction of multiple comparisons, *****p* < 0.0001, ns: not significant)

### Supplementary Tables

**Supplementary Table 1** Correlations between serum compounds and TS in 12w mice. **Supplementary Table 2** List of 12w serum SAMs between shamM and shamF under POS and NEG modes.

**Supplementary Table 3** List of 18m serum SAMs among shamM, shamF and casM under NEG and POS modes.

**Supplementary Table 4** List of 12w SAMs between shamM and casM under NEG and POS modes.

**Supplementary Table 5** Cfuzzing clustering for the 12w serum SAMs among shamM,shamF and casM.

**Supplementary Table 6** Spearman coexpression coefficient R values among the 121 CDBs. **Supplementary Table 7** BioPlanet enrichment analysis of the 51 taurine correlated genes. **Supplementary Table 8** Joint enrichment analysis between 18m shamM and shamF. **Supplementary Table 9** Joint enrichment analysis between 18m shamM and casM kidneys. **Supplementary Table 10** Joint enrichment analysis between 18m shamM and shamF livers. **Supplementary Table 11** Joint enrichment analysis between 18m shamM and casM livers.
